# Hierarchical Association Coefficient Algorithm

**DOI:** 10.1101/043844

**Authors:** Bongsong Kim

## Abstract

Suppose that members in a universal set categorized based on observations, and that categories can be stratified based on the average of observations within each category. Two sorting extremes can be obtained from the perspective of arbitrariness of an order of observations. The first sorting extreme is an increasing order of observations on ascendingly stratified categories. The second sorting extreme is a decreasing order of observations on ascendingly stratified categories. Hierarchical association coefficient (HA-coefficient) algorithm is based on a principle that any order of observations in stratified categorization can be placed between the two sorting extremes. The algorithm produces a proportion of how much an order of observations in stratified categorization is close to the first sorting extreme, or how much an order of categorized observations is distant from the second sorting extreme. This paper introduces a theory about the HA-coefficient algorithm, and shows its applications with example data. In addition, proving a reliability of the algorithm is shown through a simulation.

## Introduction

Hierarchical association coefficient algorithm (HA-coefficient algorithm) can work with observations in stratified categorization based on the average of observations within each category. The algorithm is fundamentally based on a concept that any marginal data between observations and the categorical identifiers is one outcome out of all possible permutations of the observations against categorical identifiers. From any marginal data between observations and categorical identifiers, two sorting extremes can be obtained by arranging the observations and categorical identifiers in the following manner: (1) the observations are increasingly sorted into ascendingly stratified categories; (2) the observations are decreasingly sorted into the same categories. The algorithm produces a proportion for arbitrariness of an order of observations given ascendingly stratified categorization between the two sorting extremes, and the proportion is called hierarchical association coefficient (HA-coefficient). Thus, the former and latter sorting extremes always produce the HA-coefficient = 1 and HA-coefficient = 0, respectively, so that the HA-coefficient indicates how much the observed order of observations is close to the increasing order of observations given ascendingly stratified categories. The algorithm measures the HA-coefficient in a physical manner based on distance and area.

This paper introduces the HA-coefficient algorithm, and shows its applications with three different situations. Also, it presents a demonstration that the algorithm produces a consistent result with three different simulated data sets that actually share the same pattern across the simulations. This result proves that the algorithm is reliable. The development of HA-coefficient algorithm was initially motivated to detect single nucleotide polymorphisms (SNPs) of high association with traits of interest in soybean breeding. As many disciplines need an analysis of data in stratified categorization, the HA-coefficient algorithm can be implemented to a wide spectrum of studies.

## Theory and method

### Hierarchical association distance metric

Assume that data satisfy the following conditions:

Every member has a positive real value as observation.
Every member has either nominal or ordinal identifier that determines its own category.
All categorical averages of observations are different.

Under the above conditions, the observations and categories for every member are marginal. And, multiple categories can be stratified based on the average of observations within each category. Based on these aspects, Definitions 1 to 4 were established:

#### Definition 1.

“Hierarchical” defines that categorical groups are stratified based on the average of observations within each category.

#### Definition 2.

Suppose that multiple categories are sorted in ascending order based on the average of observations within each category. “Top categorization” defines a condition that observations arranged in ascending order within each category lead to ascending order across the multiple categories.

#### Definition 3.

Suppose that multiple categories are sorted according to the top categorization according to Definition 2. “Bottom categorization” defines a condition that observations arranged in descending order within each category lead to descending order across the multiple categories.

#### Definition 4.

“Hierarchical association” defines a proportion representing how close the top and observed categorizations are, or how distant the bottom and observed categorizations are.

In Definition 1, “hierarchical” attribute is a crucial concept since all computations for the HA-coefficient algorithm require categories to be sorted in a hierarchical manner. Note that permutation of observations within a category does not affect a calculation associated with the hierarchical association. Suppose that multiple categories are hierarchically arranged in a universal set. The set can be divided into two subsets at any categorical boundary. For convenience, say that the left and right subsets were obtained, denoted by L and R, respectively. If the sum of the L subset in the top categorization is greater than that in the observed categorization, say that L is Category 1 and its sum is *x1*, and that the R is Category 2 and its sum is *x2*. Likewise, R and L can be Categories 1 and 2, respectively.

#### Definition 5.

Suppose that multiple categories and observations are marginal and hierarchically stratified in a universal set. The set can be divided into two subsets (S1 and S2) at any categorical boundary. A sum of either S1 or S2 must be greater in the top categorization than in the observed categorization, which is defined by Category 1, and its sum is defined as *x1*. Thus, the other category and its sum are defined by Category 2 and *x2*, respectively.

Definition 5 defines a rule to determine the *x1* in Category 1. As a component of HA-coefficient algorithm, the hierarchical association distance metric (HADM) calculates a *d* coefficient. Say that the *d* coefficient given a variable *x* is denoted by *d*_*x*_, where a subscript *x* on *d*_*x*_ refers to *x1* according to Definition 5. And say that *g1* and *g2* are *x1* and *x2*, respectively, in the top categorization according to Definition 5, and that *r1* and *r2* are *x1* and *x2*, respectively, in the observed categorization according to Definition Definitions 5. The *d*_*x*_ can be can be calculated as:

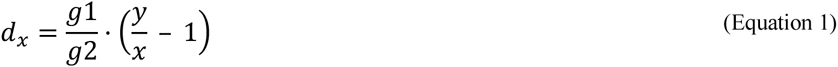

where *x* = the variable; *d*_*x*_ = the *d* coefficient given *x*; *y* = the sum of all observations; *g1* = the *x1* in the top categorization according to Definition 5; *g2* = the *x2* in the top categorization according to Definition 5.

The Equation 1 can be derived:

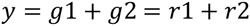

Substitute *r1* by *x* as a variable, so that *r2* = *y* − *x*. Note that *x* is a variable referring to *x1*. Then,

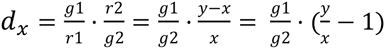

where *r1* = the *x1* in the observed categorization; *r2* = the *x2* in the observed categorization; *x* = the variable; *d*_*x*_ = the *d* coefficient given *x*; *y* = the sum of all observations; *g1* = the *x1* in the top categorization according to Definition 5; *g2* = the *x2* in the top categorization according to Definition 5.

It is always true that 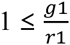 and 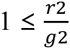, so that 1 ≤ *d*_*x*_. The top categorization according to Definition 2 always produces *d*_*x*_ = 1 as a minimum *d* coefficient, while the bottom categorization according to Definition 3 always results in maximum *d*_*x*_. The *d*_*x*_ can be interpreted as a hierarchical association distance given *x* = *x1* from any categorization. In dealing with *d*_*x*_, the categories 1 and 2 have to be hierarchical according to Definition 1. If they are not hierarchical, *d*_*x*_ is unsolvable.

### HA-coefficient algorithm with two categories

The *d*_*x*_ against *x* in the HADM (Equation 1) is shaped into a curve (see Figure 1). An area delimited between two *x1* corresponding to the bottom and top categorizations refers to cumulative hierarchical association distances from all possible permutations of the observations against marginal categorical identifiers, and say the area as W. Likewise, an area delimited between two *x1*s corresponding to the bottom and observed categorizations refers to cumulative hierarchical association distances between the two categorizations, and say the area as R. The W and R can be calculated as follows:

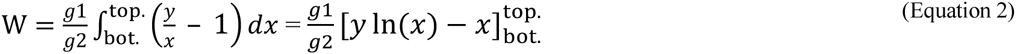

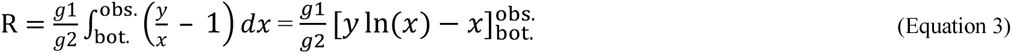

**Figure 1.**
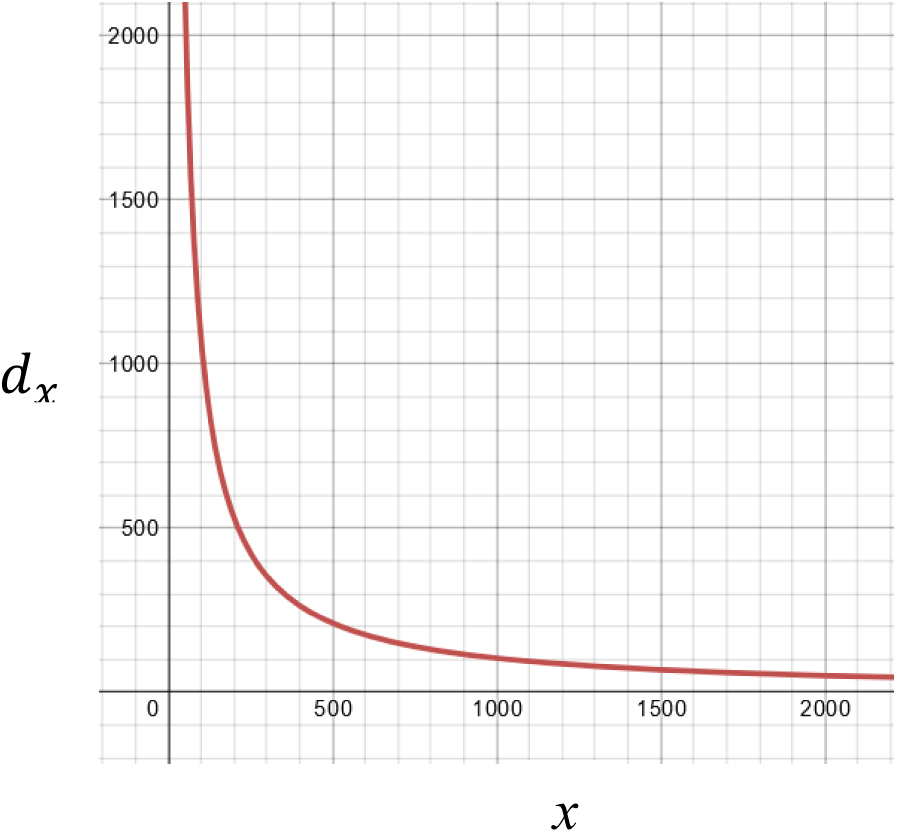
A curve for 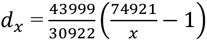. This shows a curve of *d*_*x*_ against *x* given the first boundary of SNP1 data in Table 1 (see Table 2 also), in which *x*-and *y*-axes represent scales for *x* and *d*_*x*_, respectively. The all area (W) and realized area (R) can be determined with area-in and area-out under the curve delimited. The graph was generated using online graph calculator at https://www.desmos.com/calculator.

Ultimately, the HA-coefficient can be obtained as a ratio between R and W as follows:

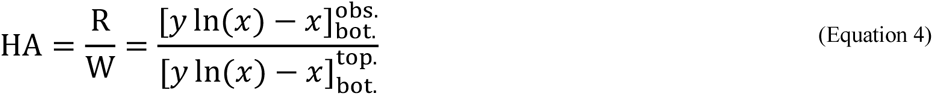

where HA = the HA-coefficient; *x* = the variable; W = the whole area that *x* can formulate between two *x1*s corresponding the bottom and top categorizations; R = the realized area that *x* can formulate between two *x1*s corresponding the bottom and observed categorizations; obs. = the *x1* in the observed categorization; top. = the *x1* in the top categorization; bot. = the *x1* in the bottom categorization; *y* = the sum of whole observations; *g1* = the *x1* in the top categorization; *g2* = the *x2* in the top categorization.

It is always true that 0 ≤ R ≤ W, so that the HA-coefficient results in a proportion. The HA-coefficient represents the hierarchical association according to Definition 4. So far, two categories of data were addressed. In reality, it is general that observations can be classified into more than two hierarchical categories. The calculation of HA-coefficient can be extended to data with any number of hierarchical categories.

### HA-coefficient algorithm with any number of categories

In case that two or more categories have equal averages, it should be carefully considered that the categories can be merged into a single category. If the categories cannot be merged due to other attribute, *d*_*x*_ is unsolvable because the categories are not hierarchical according to Definition 1. In case that a universal set nests *n* categories, the set can be classified into two subsets of *n*-1 at each categorical boundary. With multiple categories, the HA-coefficient algorithm can be implemented following two procedures: (1) computing component HA-coefficients of *n*-1 times at each categorical boundary, and (2) averaging all component HA-coefficients. As averaging method, either of harmonic and arithmetic means can be taken. The methods for computing the HA-coefficient can be denoted as follows:

HA-coefficient algorithm based on harmonic mean,

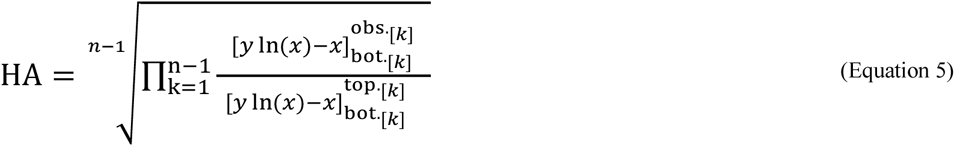
HA-coefficient algorithm based on arithmetic mean,

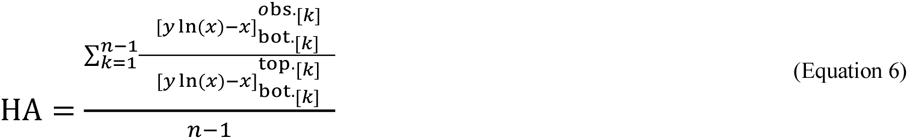

where HA = the HA-coefficient; *x* = the variable; *n* = the total number of categories; *k* = the variable referring to an order of categorical boundary; *y* = the sum of all observations; bot._[*k*]_ = the *x1* in the bottom categorization given the k’th categorical boundary; obs._[*k*]_ = the *x1* in the observed categorization given the *k*’th categorical boundary; top._[*k*]_ = the *x1* in the top categorization given the *k*’th categorical boundary.

Both Equations 5 and 6 produce identical results, and are complete forms of the HA-coefficient algorithm. In implementing the algorithms, classifying a universal set into two subsets of *n*-1 times at each categorical boundary provides the following properties:

1. computing the component HA-coefficients at each categorical boundary happens with entire data, which allows to obtain an ultimate HA-coefficient by simply averaging;
2. any change of *d*_*x*_ from permuting observations against categorical identifiers can be captured through discovering a change of a ratio between *x1* and *x2*.

### Independent categories and dependent categories

The categorization can be hierarchical in pre-determined (pre-hierarchical) or post-determined (post-hierarchical) manner. If categories are determined independent from the observations, categories are pre-hierarchical. If categories are determined dependent on the observations, categories are post-hierarchical.

#### Definition 6.

When categorical hierarchy is determined independent from observations, categories are “prehierarchical”. When categorical hierarchy is determined dependent on observations, categories are “post-hierarchical”.

It is important to note that Definition 6 determines a feasibility that HA-coefficient = 0. The independent categorization makes it feasible that HA-coefficient = 0, while the dependent categorization does not make it feasible that HA-coefficient = 0. This will be addressed in detail across the three example situations in Results and Discussion.

### Curve of *d*_*x*_

Figure 1 shows a part of curve for 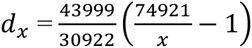, which is derived from SNP1 scores given soybean yield quantity at the boundary between M0 and (M1, M2) in Table 1 (see Table 2 also). On the graph, *x*-axis and *y*-axis represent a scale of *x* and *d*_*x*_, respectively. The *d*_*x*_ curve can be used to determine both the W and R with setups of both area-in and area-out. On the graph, the *x* can always have a value between two *x1*s responding to the bottom and top categorizations. Note that the *x* can also be equal to either of *x1*s responding to the bottom and top categorizations, which will result in HA-coefficient = 0 or HA-coefficient = 1, respectively. As shown in Figure 1, the *d*_*x*_ curve can be drawn only in quadrant I, which indicates that only positive real numbers are available as observations.

**Table 1.** Simulated SNPs’ scores and yield quantity (kg / ha) for 20 soybean varieties. At the bottom of table, rounded means within each category are summarized, and its hierarchical ranking is shown in parentheses.

| ID | SNP1 | SNP2 | SNP3 | Yield (kg/ha) |
| --- | --- | --- | --- | --- |
| Soybean.1 | 0 | 0 | 1 | 3636 |
| Soybean.2 | 1 | 1 | 1 | 3866 |
| Soybean.3 | 0 | 0 | 2 | 4012 |
| Soybean.4 | 0 | 1 | 2 | 3583 |
| Soybean.5 | 1 | 1 | 0 | 4075 |
| Soybean.6 | 2 | 2 | 2 | 3957 |
| Soybean.7 | 1 | 0 | 2 | 4054 |
| Soybean.8 | 2 | 2 | 1 | 3927 |
| Soybean.9 | 0 | 2 | 2 | 3255 |
| Soybean.10 | 2 | 1 | 2 | 3994 |
| Soybean.11 | 2 | 0 | 2 | 3904 |
| Soybean.12 | 0 | 1 | 0 | 3927 |
| Soybean.13 | 2 | 2 | 0 | 3603 |
| Soybean.14 | 0 | 0 | 0 | 4196 |
| Soybean.15 | 0 | 1 | 2 | 3573 |
| Soybean.16 | 1 | 2 | 2 | 3082 |
| Soybean.17 | 0 | 2 | 0 | 2711 |
| Soybean.18 | 0 | 0 | 0 | 3613 |
| Soybean.19 | 2 | 0 | 1 | 3955 |
| Soybean.20 | 1 | 0 | 2 | 3998 |
| Mean 0 (M0) | 3612 (1) | 3921 (3) | 3688 (1) | Total<br>74921 |
| Mean 1 (M1) | 3815 (2) | 3836 (2) | 3846 (3) |  |
| Mean 2 (M2) | 3890 (3) | 3423 (1) | 3741 (2) |  |

**Table 2.** Summary of *x1* and *x2* extracted from Table 1. The *x1* and *x2* were determined at each categorical boundary according to Definition 5. By applying the HA-coefficient algorithm to the *x1*s, the HA-coefficient can be obtained. Hierarchical ranking of categories is ascendingly present within parentheses underneath each SNP name in the first column. The second column shows classifications of the whole set using a vertical bar and parentheses, and *x1* and *x2* are marked within square brackets. As all the three SNPs have three categories, in common, the whole set can be classified at two different categorical boundaries at a time for each SNP.

| SNP | Classification of $x1$ and $x2$ | Category type | $x1$ | $x2$ | Total |
| --- | --- | --- | --- | --- | --- |
| SNP1<br>(M0, M1, M2) | M0 \| (M1, M2)<br>[$x2$ \| $x1$] | Observed | 42415 | 32506 | 74921 |
|  |  | Top | 43999 | 30922 | 74921 |
|  |  | Bottom | 38753 | 36168 | 74921 |
| | (M0, M1) \| M2<br>[$x2$ \| $x1$] | Observed | 23340 | 51581 | 74921 |
|  |  | Top | 24329 | 50592 | 74921 |
|  |  | Bottom | 19807 | 55114 | 74921 |
| SNP2<br>(M2, M1, M0) | M2 \| (M1, M0)<br>[$x2$ \| $x1$] | Observed | 54386 | 20535 | 74921 |
|  |  | Top | 55114 | 19807 | 74921 |
|  |  | Bottom | 50592 | 24329 | 74921 |
| | (M2, M1) \| M0<br>[$x2$ \| $x1$] | Observed | 31368 | 43553 | 74921 |
|  |  | Top | 32241 | 42680 | 74921 |
|  |  | Bottom | 27056 | 47865 | 74921 |
| SNP3<br>(M0 M2, M1) | M0 \| (M2, M1)<br>[$x2$ \| $x1$] | Observed | 52796 | 22125 | 74921 |
|  |  | Top | 55114 | 19807 | 74921 |
|  |  | Bottom | 50592 | 24329 | 74921 |
| | (M0, M2) \| M1<br>[$x2$ \| $x1$] | Observed | 15384 | 59537 | 74921 |
|  |  | Top | 16337 | 58584 | 74921 |
|  |  | Bottom | 12621 | 62300 | 74921 |

### Data generation for simulations

Four simulated data sets were generated using R package (R Core Team, 2014). Of these, three were used to show an application of practice the HA-coefficient algorithm, and the other three were one was used to prove a reliability of the algorithm. The first three data former sets were manually adjusted after generation.

### Simulations to prove a reliability of HA-coefficient algorithm

In order to see if the HA-coefficient algorithm is robust, three simulated data sets in simple design were generated. The Respective simulated data sets have two to four categories. In the simulation, the following two questions were focused on, which are:

1. Do the resulting HA-coefficients gradually increase as categorizations change from arbitrary to top?
2. Does the HA-coefficient algorithm produce the same result for three different shapes of data that are inherently in the same pattern?

As an example, the simulated data set where three categorical identifiers and observations are marginal was made in the following steps:

1. Create a vector consisted of 1,200 numbers from 3 to 3,600 at an interval of 3 in ascending order. These values will be used as observations.
2. Create a 1,200 by 1,200 matrix filled with categorical identifiers, 0, 1, and 2 generated randomly.
3. Divide the matrix equally into three areas, vertically. Subsequently, overwrite 0, 1, and 2 on the area above the diagonal at each of the third, respectively.
4. Append the vector for the observations to the right end of the matrix. By doing so, the matrix size becomes 1,200 by 1,201.
5. Create a repository vector that will contain the 1,200 resulting HA-coefficients.
6. Calculate the HA-coefficient with a pair of columns for identifiers and observations across the matrix of identifiers.
7. Append the resulting HA-coefficients to the repository vector.
8. Repeat the above steps 100 times, and average the 100 repository vectors.

Above procedures illustrate generating the simulated data that have three categories. The simulated data carrying the two and four hierarchical categories were prepared in a manner of dividing the entire matrix into two and four sub-areas, respectively. The simulated data sets can be seen in Figure 2. The designs of the simulations intend to output gradually increasing HA-coefficients as a column of the matrix changes from left to right. In order to draw a smooth plot, the entire cycle of the simulation was repeated 100 times, and a plot was obtained through averaging.

**Figure 2.**
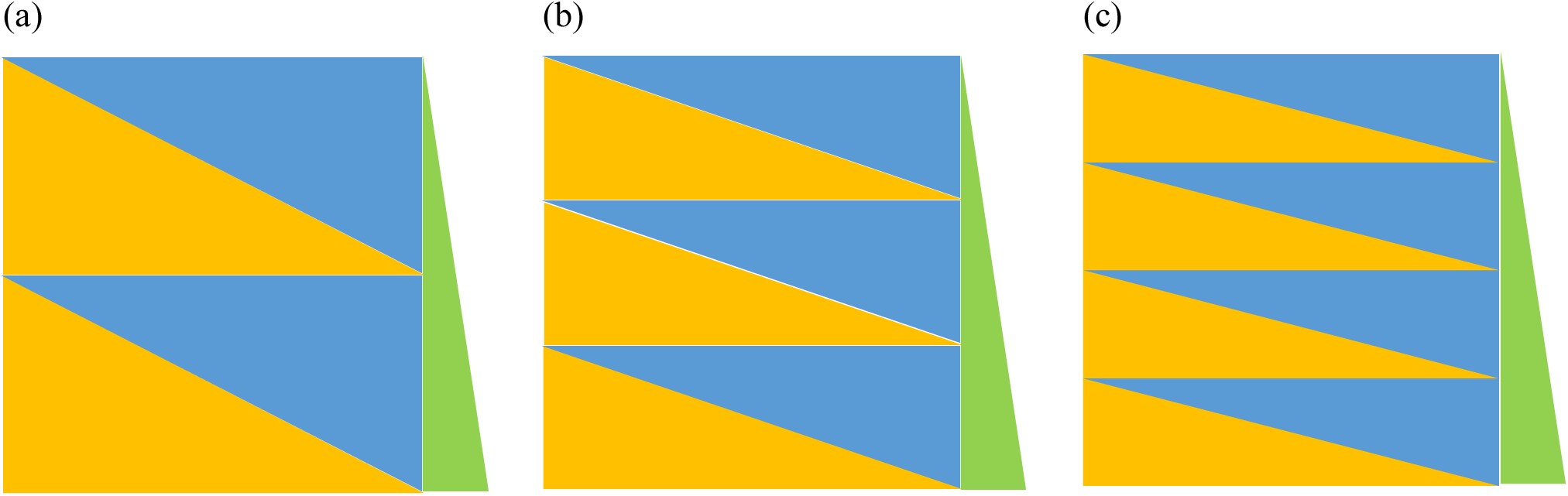
Three simulated data sets. (a), (b), and (c) are filled with categorical identifiers of two-, three-, and four-types, respectively. Each square includes multiple blue triangles. Within each square, the same identifiers are filled in each blue triangle, but different blue triangles have different identifiers. Yellow triangle contains arbitrary categorical identifiers, and green triangle on the right side refers to a vector containing 1,200 observations from 3 to 3,600 at an interval of 3. When a pair of columns for the left end of the matrix and observations are marginal, arbitrary categorization will be obtained, while the top categorization will be formed with a pair of columns for 1,200^th^ column and observations.

## Results and discussion

Let us apply the HA-coefficient algorithm to the following three example situations. The three examples refer to varying situations based on a dependency between categories and observations according to Definition 6. Situation 1 presents that categories are dependent on observations, while Situations 2 and 3 show that categories are independent from observations. Situation 3 is an example that categories are determined by multi-dimensional observations.

### Situation 1: Categories fully depending on observations

Table 1 displays simulated genotypes and phenotypes, which are categorized based on singlenucleotide polymorphisms (SNPs) and observations for yield (kg/ha), respectively. The SNPs classify 20 soybean varieties into three categories (0, 1, and 2), and a hierarchical level among categories can be determined based on the average of yield within each category. This is a typical post-hierarchical case according to Definition 6 since the hierarchical categories are obtained depending on observations. This condition makes it impossible to arrange an order of observations to be in the bottom categorization since the hierarchical ranking of the categories can be altered with permuting the observations against categorical identifiers. Once categories are fixed based on the observations, the bottom and top categories can be determined. The scores from each SNP were resorted according to the top and bottom categorizations, which are tabulated in Supplementary table 1. By referring to the resorted observations, the *x1*s and *x2*s for each SNP can be determined according to Definition 5 at each categorical boundary. Table 2 shows a summary for both *x1*s and *x2*s for the three SNPs in Table 1. Basically, Table 2 shows a division of the total sum into two sums at each categorical boundary. Therefore, total sums are consistent across all boundaries, but divided sums vary depending on classification.

By applying Equation 5 to the *x1*s from Table 2, the HA-coefficients for the three SNPs were calculated as follows:

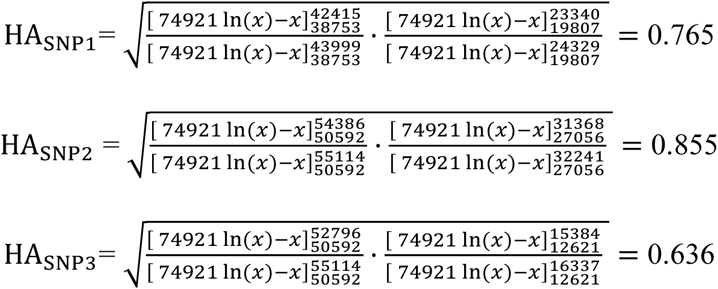

where HA_SNP1_= the HA-coefficient for SNP1; HA_SNP2_= the HA-coefficient for SNP2; HA_SNP3_= the HA-coefficient for SNP3; *x* = the variable.

From the above results, it was identified that SNP2 has the strongest hierarchical association, followed by SNP1 and SNP3.

### Situation 2: Categories independent from observations

When hierarchical ranking of categories is given from an external source, it satisfies a pre-hierarchical condition that categories and observations are independent according to Definition 6. Table 3 shows the scores from two Mathematics quizzes for the same 27 students. Based on the scores from Quiz 1, a teacher classifies all students into three hierarchical categories upon the following criteria:

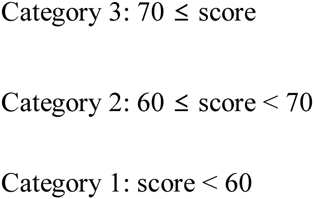

**Table 3.** Simulated scores of two Mathematics quizzes for 27 students. The three categories were classified based on the scores from Quiz 1, in which Categories 3, 2, and 1 satisfy conditions: 70 ≤ score, 60 ≤ score < 70, and score < 60, respectively.

| Category | ID | Mathematics score |  |
| --- | --- | --- | --- |
|  |  | Quiz 1 | Quiz 2 |
| Category 3<br>(C3) | Student.1 | 88 | 79 |
|  | Student.2 | 83 | 82 |
|  | Student.3 | 81 | 80 |
|  | Student.4 | 87 | 73 |
|  | Student.5 | 78 | 70 |
|  | Student.6 | 71 | 52 |
|  | Student.7 | 71 | 75 |
|  | Student.8 | 75 | 61 |
| Category 2<br>(C2) | Student.9 | 69 | 57 |
|  | Student.10 | 68 | 77 |
|  | Student.11 | 68 | 67 |
|  | Student.12 | 67 | 59 |
|  | Student.13 | 66 | 54 |
|  | Student.14 | 66 | 46 |
|  | Student.15 | 64 | 66 |
|  | Student.16 | 62 | 53 |
|  | Student.17 | 61 | 85 |
| Category 1<br>(C1) | Student.18 | 58 | 68 |
|  | Student.19 | 58 | 69 |
|  | Student.20 | 57 | 51 |
|  | Student.21 | 56 | 61 |
|  | Student.22 | 55 | 72 |
|  | Student.23 | 52 | 61 |
|  | Student.24 | 52 | 67 |
|  | Student.25 | 48 | 75 |
|  | Student.26 | 50 | 68 |
|  | Student.27 | 49 | 43 |

If the ranking of scores from Quiz 2 is preserved or exchanged within each category based on Quiz 1, there will be no difference on hierarchical association between results from the two quizzes. However, new scores from different quiz for the same students are often differently ranked across categories, so that the teacher could raise the following questions. Was the categorical stratification based on Quiz 1 maintained in the result from Quiz 2? If not, how much of categorical solidity based on Quiz 1 remains in the result from Quiz 2? In order for the HA-coefficient algorithm to answer this question, the categorization determined based on Quiz 1 has to be fixed, and applied to the result from Quiz 2. The scores from Quiz 2 were resorted according to the top and bottom categorizations, which are tabulated in Supplementary table 2. Table 4 presents the *x1*s and *x2*s at each categorical boundary. The HA-coefficient were calculated by applying Equation 5 to the *x1*s from Table 4 as follows:

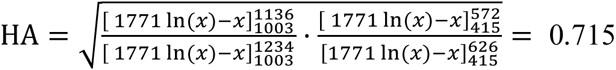

where HA = the HA-coefficient; *x* = the variable.

**Table 4.** Summary of *x1*s and *x2*s extracted from Table 3. The first column shows hierarchical ranking of three categories (C1, C2, and C3) and classified sets (*x1* and *x2*), in which a vertical bar is a boundary to classify the whole set, and parentheses collapse categories into the same set.

| Classification of<br>$x1$ and $x2$ | Category<br>type | $x1$ | $x2$ | Total |
| --- | --- | --- | --- | --- |
| C1 \| (C2, C3)<br>[$x2$ \| $x1$] | Observed | 1136 | 635 | 1771 |
|  | Top | 1234 | 537 | 1771 |
|  | Bottom | 1003 | 768 | 1771 |
| (C1, C2) \| C3<br>[$x2$ \| $x1$] | Observed | 572 | 1199 | 1771 |
|  | Top | 626 | 1145 | 1771 |
|  | Bottom | 415 | 1356 | 1771 |

The above result indicates that the result from Quiz 2 keeps 71.5 % of solidity of categorization determined by Quiz 1.

### Situation 3: Categories determined by multi-dimensional observations

It is possible that multi-dimensional observations form a categorization. In this case, each dimension of observations influences the categorization. Table 5 displays simulated data about results from Mathematics and English quizzes. The table includes Mathematics scores, English scores, and average for the same 27 students, and the students are hierarchically categorized based on the average. Thus, the Mathematics and English scores are related to the categorization. The criteria for the categorization are as follows:

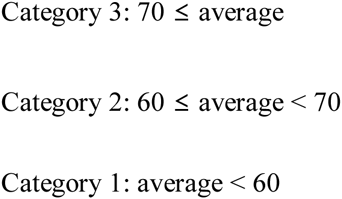

**Table 5.** Simulated scores of Mathematics and English quizzes for 27 students. The three categories were classified based on averages, in which Categories 3, 2, and 1 (C3, C2, and C1) satisfy conditions: 70 ≤ average score, 60 ≤ average score < 70, and average score < 60, respectively.

| Category | ID | Score |  | Average |
| --- | --- | --- | --- | --- |
|  |  | Mathematics | English |  |
| Category 3<br>(C3) | Student.1 | 88 | 79 | 83.5 |
|  | Student.2 | 83 | 82 | 82.5 |
|  | Student.3 | 87 | 73 | 80.0 |
|  | Student.4 | 68 | 90 | 79.0 |
|  | Student.5 | 78 | 70 | 74.0 |
|  | Student.6 | 81 | 66 | 73.5 |
|  | Student.7 | 61 | 85 | 73.0 |
|  | Student.8 | 71 | 75 | 73.0 |
|  | Student.9 | 68 | 77 | 72.5 |
| Category 2<br>(C2) | Student.10 | 58 | 81 | 69.5 |
|  | Student.11 | 71 | 67 | 69.0 |
|  | Student.12 | 67 | 71 | 69.0 |
|  | Student.13 | 69 | 66 | 67.5 |
|  | Student.14 | 55 | 72 | 63.5 |
|  | Student.15 | 58 | 68 | 63.0 |
|  | Student.16 | 75 | 50 | 62.5 |
|  | Student.17 | 48 | 75 | 61.5 |
|  | Student.18 | 64 | 57 | 60.5 |
|  | Student.19 | 66 | 54 | 60.0 |
| Category 1<br>(C1) | Student.20 | 52 | 67 | 59.5 |
|  | Student.21 | 50 | 68 | 59.0 |
|  | Student.22 | 56 | 61 | 58.5 |
|  | Student.23 | 62 | 53 | 57.5 |
|  | Student.24 | 52 | 61 | 56.5 |
|  | Student.25 | 66 | 46 | 56.0 |
|  | Student.26 | 57 | 51 | 54.0 |
|  | Student.27 | 49 | 43 | 46.0 |

The size of each category and categorical classification of each student are determined based on the averages, so that this situation is pre-hierarchical. If the ranking from each quiz is preserved or exchanged within each category based on the averages, there will be no difference on hierarchical association between each quiz and average. However, the scores from the two quizzes ranked across categories, so that the following questions could be raised. Was the categorical stratification based on the averages maintained in the result from each quiz? If not, how much of categorical solidity based on the averages remains in the result from each quiz? To answer the questions using the HA-coefficient algorithm, the given categorization has to be fixed, and applied to the scores from each quiz. The scores were resorted according to the top and bottom categorizations, which are tabulated in Supplementary table 3. Table 6 shows the *x1*s and *x2*s at each categorical boundary. By applying Equation 5 to the *x1*s from Table 6, the HA-coefficients were calculated as follows:

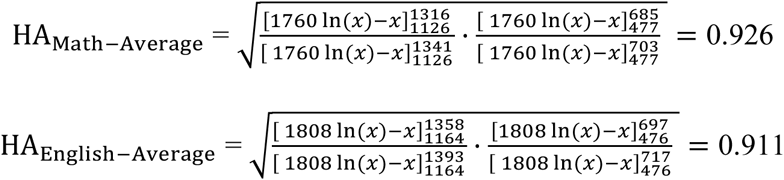

where HA_Math‒Average_ = the HA-coefficient for the Mathematics scores given the averages; HA_English‒Average_= the HA-coefficient for the English scores the given averages; *x* = the variable.

**Table 6.** Summary of both *x1* and *x2* extracted from Table 5. The second column shows hierarchical ranking of the three categories (C1, C2, and C3) and sets classified into *x1* and *x2*, in which a vertical bar is a boundary to classify the whole set, and parentheses collapse two categories into the same set.

| Subject | Classification of $x1$ and $x2$ | Category type | $x1$ | $x2$ | Total |
| --- | --- | --- | --- | --- | --- |
| Mathematics | C1 \| (C2,C3)<br>[$x2$ \| $x1$] | Observed | 1316 | 444 | 1760 |
|  |  | Top | 1341 | 419 | 1760 |
|  |  | Bottom | 1126 | 634 | 1760 |
| | (C1,C2) \| C3<br>[$x2$ \| $x1$] | Observed | 685 | 1075 | 1760 |
|  |  | Top | 703 | 1057 | 1760 |
|  |  | Bottom | 477 | 1283 | 1760 |
| English | C1 \| (C2,C3)<br>[$x2$ \| $x1$] | Observed | 1358 | 450 | 1808 |
|  |  | Top | 1393 | 415 | 1808 |
|  |  | Bottom | 1164 | 644 | 1808 |
| | (C1,C2) \| C3<br>[$x2$ \| $x1$] | Observed | 697 | 1111 | 1808 |
|  |  | Top | 717 | 1091 | 1808 |
|  |  | Bottom | 476 | 1332 | 1808 |

The above results explain that the Mathematics and English scores keep 92.6 and 91.1 % of solidities of the categorization determined by the averages, respectively.

### Simulation to evaluate a reliability of HA-coefficient algorithm

In order to verify a reliability of the algorithm, the simulation was conducted with the two following questions:

1. As a categorization is getting from arbitrary to top, does the algorithm produce an increasing HA-coefficient?
2. Does the HA-coefficient algorithm produce the same result for three different shapes of data that are inherently in the same pattern?

The above questions are very essential to evaluate the reliability of the algorithm. For the simulation, three data sets in square dimension (1,200 by 1,200) were generated. The first, second and third squares have two, three and four categorical identifiers within each column, respectively. Each simulated data set was vertically partitioned into two to four areas with equal size. And, a vector containing 1,200 numbers from 3 to 3,600 at regular interval of 3 was appended at the right end of each matrix. Figure 2 visualizes the structures of the simulated data. Within each square, each blue triangle is filled with the same categorical identifiers, but different blue triangles have different categorical identifiers. Yellow triangles refer to area for arbitrarily mixed categorical identifiers. Meanwhile, each figure has a green triangle with a slope, which implies an increasing pattern of observations. Thus, when pairing two columns for categorical identifiers and observations, an arrangement of categorical identifiers given the observations gradually transpositions from arbitrary to top as the column changes from left to right. A relationship between the categories and observations stays dependent to some degree of right column from the left end, but shifts to solid independent condition since averages for each category are gradually getting hierarchically solid as a column changes from left to right. The implementation of the HA-coefficient algorithm was repeated 100 times with each simulated data to minimize a noisy fluctuation. Figure 3 shows three outcome plots, illustrating a steady increasing pattern. This returns a positive answer to the first question. Meanwhile, the shapes of the three plots in Figure 3 seemingly look closely similar, and each plot ranges between around 0.75 and 1. Since the categorical identifiers and observations are post-hierarchical from the first column to some degree of depth to the right, these simulations do not allow to determine the bottom categorization same as explained in Situation 1. Therefore, it cannot happen that HA-coefficient = 0 with these simulations. Importantly, the three different simulations share two following properties: (1) equal proportions of identifiers according to the top categorization in column-wise comparison; (2) the identical observations in the same order. These two properties assure that the three different simulated data have the same pattern in a fashion of column-wise comparison. Therefore, in order for the HA-coefficient algorithm to be robust, the same results have to be obtained from the three simulations. Table 7 shows that the Pearson correlation coefficients among the three plots are remarkably close to one another. This returns a positive answer to the second question, and indicate that the averaging with *n*-1, where *n* is the number of categories, does not perturb calculating the HA-coefficient.

**Figure 3.**
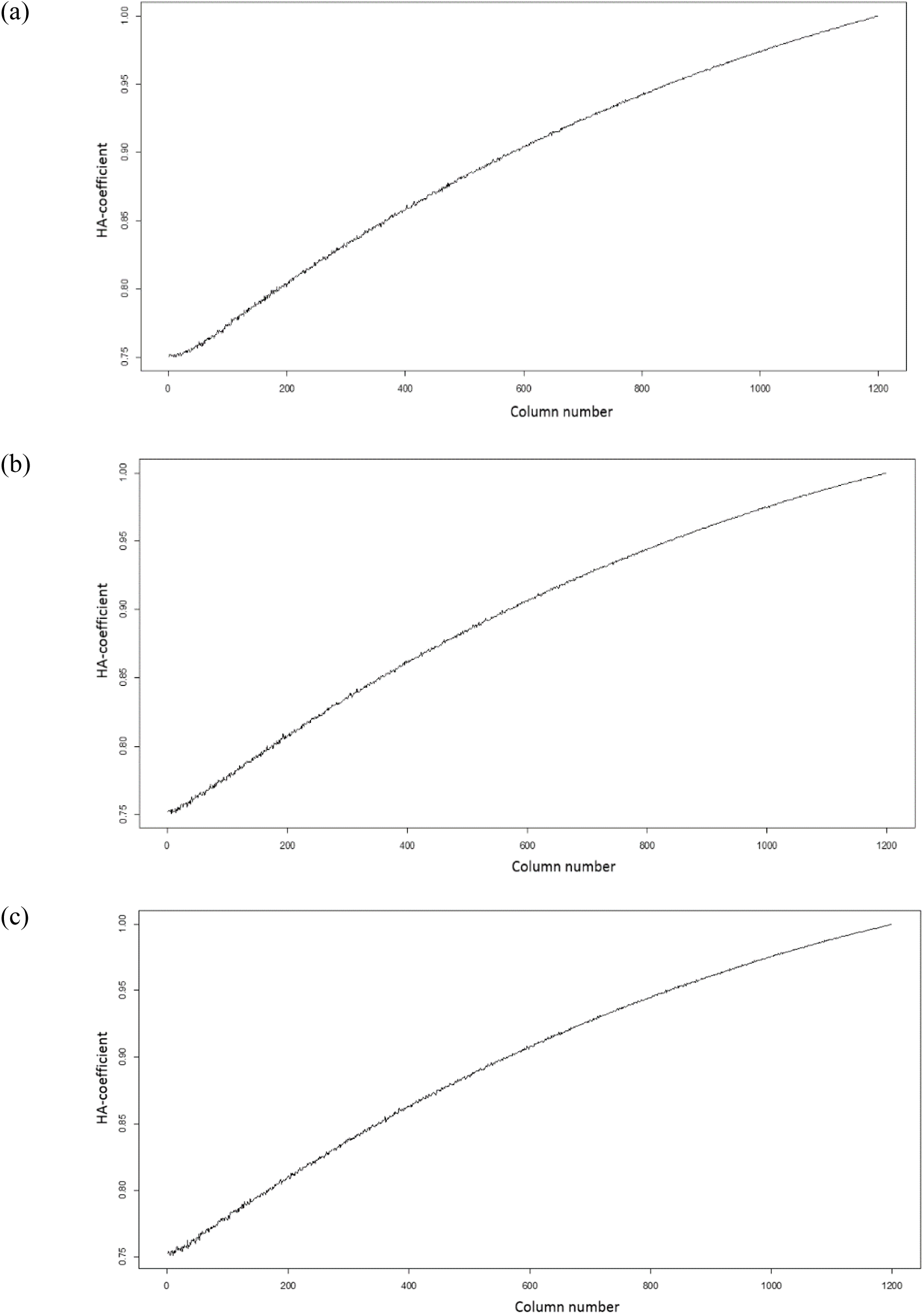
Plots (a), (b), and (c) are drawn based on the HA-coefficients resulted from the simulated data in Figure 2 (a), (b), and (c), respectively. Each plot was obtained through averaging 100 repetitions of implementing the HA-coefficient algorithm.

**Table 7.** Pearson correlation coefficients among the three plots in Figure 3

| Number of categories | 2 | 3 | 4 |
| --- | --- | --- | --- |
| 2 | 1 | 0.999893 | 0.999845 |
| 3 | 0.999893 | 1 | 0.999883 |
| 4 | 0.999845 | 0.999883 | 1 |

## Conclusion

Above, it was proven that the HA-coefficient algorithm returns a reliable result through the simulations. As long as the categories and observations are marginal, and categories are hierarchically stratified, the algorithm solves a reasonable proportion for HA-coefficient from data. It is important to note that the HA-coefficient is an objective measure rather than a statistical inference. The algorithm is easy to computerize, and its implementation does not require high computing power. The HA-coefficient algorithm was applied to real agricultural data including 4,312 SNPs and yield quantity for 5,180 soybean lines, its computation was done in 34 seconds on a laptop (Dell E6540) using R. This validates that the algorithm runs very fast on an ordinary computer. As many disciplines deal with data in hierarchical stratification, the algorithm will be useful for a wide spectrum of studies.

## Acknowledgement

Funding provided by North Central Region Soybean Program, United Soybean Board, and Department of Agronomy, Iowa State University.

---

**Supplementary table 1.**
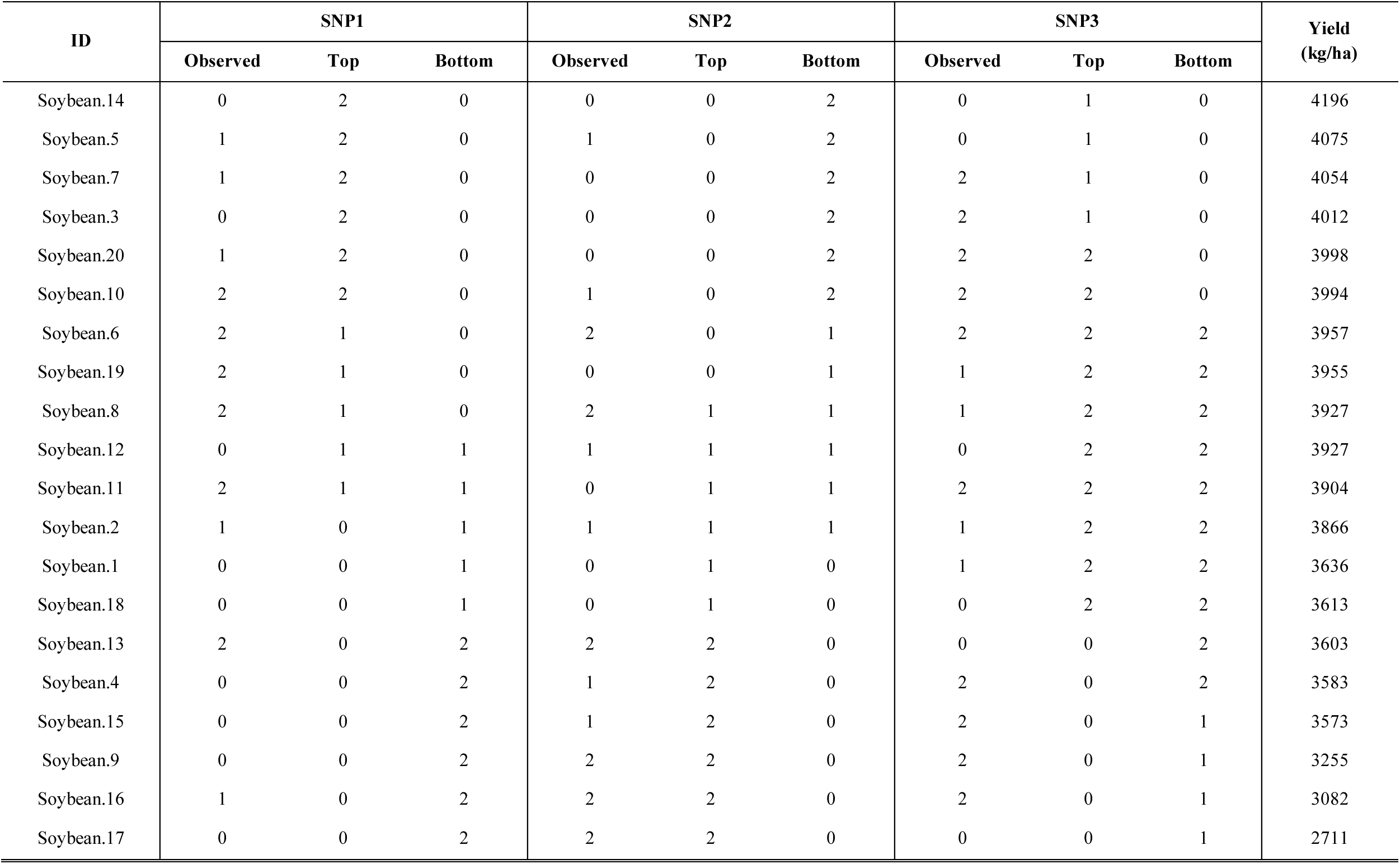
Simulated data displaying arrangements of SNP scores according to the observed, top, and bottom categorizations against decreasingly sorted yield quantity (kg/ha).

| ID | SNP1 |  |  | SNP2 |  |  | SNP3 |  |  | Yield<br>(kg/ha) |
| --- | --- | --- | --- | --- | --- | --- | --- | --- | --- | --- |
|  | Observed | Top | Bottom | Observed | Top | Bottom | Observed | Top | Bottom |  |
| Soybean.14 | 0 | 2 | 0 | 0 | 0 | 2 | 0 | 1 | 0 | 4196 |
| Soybean.5 | 1 | 2 | 0 | 1 | 0 | 2 | 0 | 1 | 0 | 4075 |
| Soybean.7 | 1 | 2 | 0 | 0 | 0 | 2 | 2 | 1 | 0 | 4054 |
| Soybean.3 | 0 | 2 | 0 | 0 | 0 | 2 | 2 | 1 | 0 | 4012 |
| Soybean.20 | 1 | 2 | 0 | 0 | 0 | 2 | 2 | 2 | 0 | 3998 |
| Soybean.10 | 2 | 2 | 0 | 1 | 0 | 2 | 2 | 2 | 0 | 3994 |
| Soybean.6 | 2 | 1 | 0 | 2 | 0 | 1 | 2 | 2 | 2 | 3957 |
| Soybean.19 | 2 | 1 | 0 | 0 | 0 | 1 | 1 | 2 | 2 | 3955 |
| Soybean.8 | 2 | 1 | 0 | 2 | 1 | 1 | 1 | 2 | 2 | 3927 |
| Soybean.12 | 0 | 1 | 1 | 1 | 1 | 1 | 0 | 2 | 2 | 3927 |
| Soybean.11 | 2 | 1 | 1 | 0 | 1 | 1 | 2 | 2 | 2 | 3904 |
| Soybean.2 | 1 | 0 | 1 | 1 | 1 | 1 | 1 | 2 | 2 | 3866 |
| Soybean.1 | 0 | 0 | 1 | 0 | 1 | 0 | 1 | 2 | 2 | 3636 |
| Soybean.18 | 0 | 0 | 1 | 0 | 1 | 0 | 0 | 2 | 2 | 3613 |
| Soybean.13 | 2 | 0 | 2 | 2 | 2 | 0 | 0 | 0 | 2 | 3603 |
| Soybean.4 | 0 | 0 | 2 | 1 | 2 | 0 | 2 | 0 | 2 | 3583 |
| Soybean.15 | 0 | 0 | 2 | 1 | 2 | 0 | 2 | 0 | 1 | 3573 |
| Soybean.9 | 0 | 0 | 2 | 2 | 2 | 0 | 2 | 0 | 1 | 3255 |
| Soybean.16 | 1 | 0 | 2 | 2 | 2 | 0 | 2 | 0 | 1 | 3082 |
| Soybean.17 | 0 | 0 | 2 | 2 | 2 | 0 | 0 | 0 | 1 | 2711 |

**Supplementary table 2.** Resorted scores from the Mathematics Quiz 2 according to the observed, top, and bottom categorizations. Students are hierarchically categorized based on Quiz 1 according to the following criteria: 70 ≤ average, 60 ≤ average < 70, and average < 60.

| Category | ID | Mathematics score |  |  |  |
| --- | --- | --- | --- | --- | --- |
|  |  | Quiz 1 | Quiz 2 |  |  |
|  |  |  | Observed | Top | Bottom |
| Category 3<br>(C3) | Student.1 | 88 | 79 | 85 | 43 |
|  | Student.2 | 83 | 82 | 82 | 46 |
|  | Student.3 | 81 | 80 | 80 | 51 |
|  | Student.4 | 87 | 73 | 79 | 52 |
|  | Student.5 | 78 | 70 | 77 | 53 |
|  | Student.6 | 71 | 52 | 75 | 54 |
|  | Student.7 | 71 | 75 | 75 | 57 |
|  | Student.8 | 75 | 61 | 73 | 59 |
| Category 2<br>(C2) | Student.9 | 69 | 57 | 72 | 61 |
|  | Student.10 | 68 | 77 | 70 | 61 |
|  | Student.11 | 68 | 67 | 69 | 61 |
|  | Student.12 | 67 | 59 | 68 | 66 |
|  | Student.13 | 66 | 54 | 68 | 67 |
|  | Student.14 | 66 | 46 | 67 | 67 |
|  | Student.15 | 64 | 66 | 67 | 68 |
|  | Student.16 | 62 | 53 | 66 | 68 |
|  | Student.17 | 61 | 85 | 61 | 69 |
| Category 1<br>(C1) | Student.18 | 58 | 68 | 61 | 70 |
|  | Student.19 | 58 | 69 | 61 | 72 |
|  | Student.20 | 57 | 51 | 59 | 73 |
|  | Student.21 | 56 | 61 | 57 | 75 |
|  | Student.22 | 55 | 72 | 54 | 75 |
|  | Student.23 | 52 | 61 | 53 | 77 |
|  | Student.24 | 52 | 67 | 52 | 79 |
|  | Student.25 | 48 | 75 | 51 | 80 |
|  | Student.26 | 50 | 68 | 46 | 82 |
|  | Student.27 | 49 | 43 | 43 | 85 |

**Supplementary table 3.** Resorted scores from the Mathematics and English quizzes according to the observed, top, and bottom categorizations. Students are hierarchically categorized based on averages according to the following criteria: 70 ≤ score, 60 ≤ score < 70, and score < 60.

| Category | ID | Mathematics score |  |  | English score |  |  | Average |
| --- | --- | --- | --- | --- | --- | --- | --- | --- |
|  |  | Observed | Top | Bottom | Observed | Top | Bottom |  |
| Category 3<br>(C3) | Student.1 | 88 | 88 | 48 | 79 | 90 | 43 | 83.5 |
|  | Student.2 | 83 | 87 | 49 | 82 | 85 | 46 | 82.5 |
|  | Student.3 | 87 | 83 | 50 | 73 | 82 | 50 | 80.0 |
|  | Student.4 | 68 | 81 | 52 | 90 | 81 | 51 | 79.0 |
|  | Student.5 | 78 | 78 | 52 | 70 | 79 | 53 | 74.0 |
|  | Student.6 | 81 | 75 | 55 | 66 | 77 | 54 | 73.5 |
|  | Student.7 | 61 | 71 | 56 | 85 | 75 | 57 | 73.0 |
|  | Student.8 | 71 | 71 | 57 | 75 | 75 | 61 | 73.0 |
|  | Student.9 | 68 | 69 | 58 | 77 | 73 | 61 | 72.5 |
| Category 2<br>(C2) | Student.10 | 58 | 68 | 58 | 81 | 72 | 66 | 69.5 |
|  | Student.11 | 71 | 68 | 61 | 67 | 71 | 66 | 69.0 |
|  | Student.12 | 67 | 67 | 62 | 71 | 70 | 67 | 69.0 |
|  | Student.13 | 69 | 66 | 64 | 66 | 68 | 67 | 67.5 |
|  | Student.14 | 55 | 66 | 66 | 72 | 68 | 68 | 63.5 |
|  | Student.15 | 58 | 64 | 66 | 68 | 67 | 68 | 63.0 |
|  | Student.16 | 75 | 62 | 67 | 50 | 67 | 70 | 62.5 |
|  | Student.17 | 48 | 61 | 68 | 75 | 66 | 71 | 61.5 |
|  | Student.18 | 64 | 58 | 68 | 57 | 66 | 72 | 60.5 |
|  | Student.19 | 66 | 58 | 69 | 54 | 61 | 73 | 60.0 |
| Category 1<br>(C1) | Student.20 | 52 | 57 | 71 | 67 | 61 | 75 | 59.5 |
|  | Student.21 | 50 | 56 | 71 | 68 | 57 | 75 | 59.0 |
|  | Student.22 | 56 | 55 | 75 | 61 | 54 | 77 | 58.5 |
|  | Student.23 | 62 | 52 | 78 | 53 | 53 | 79 | 57.5 |
|  | Student.24 | 52 | 52 | 81 | 61 | 51 | 81 | 56.5 |
|  | Student.25 | 66 | 50 | 83 | 46 | 50 | 82 | 56.0 |
|  | Student.26 | 57 | 49 | 87 | 51 | 46 | 85 | 54.0 |
|  | Student.27 | 49 | 48 | 88 | 43 | 43 | 90 | 46.0 |

